# Endodermal differentiation is reconstructed by coordination of two parallel signaling systems derived from the stele in roots

**DOI:** 10.1101/210872

**Authors:** Pengxue Li, Qiaozhi Yu, Chunmiao Xu, Xu Gu, Shilian Qi, Hong Wang, Fenglin Zhong, Shuang Wu

**Author notes:** These authors contribute equally (Pengxue Li, Qiaozhi Yu).

## Abstract

The plant roots represent the exquisitely controlled cell fate map in which different cell types undergo a complete status transition from stem cell division and initial fate specification, to the terminal differentiation. The endodermis is initially specified in meristem but further differentiates to form Casparian strips (CSs), the apoplastic barrier in the mature zone for the selective transport between stele and outer tissues, and thus is regarded as plant inner skin. In the Arabidopsis thaliana root the transcription factors SHORTROOT (SHR) regulate asymmetric cell division in cortical initials to separate endodermal and cortex cell layer. In this paper, we utilized synthetic approach to examine the reconstruction of fully functional Casparian strips in plant roots. Our results revealed that SHR serves as a master regulator of a hierarchical signaling cascade that, combined with stele-derived small peptides, is sufficient to rebuild the functional CS in non-endodermal cells. This is a demonstration of the deployment of two parallel signaling systems, in which both apoplastic and symplastic communication were employed, for coordinately specifying the endodermal cell fate.

## Introduction

Plant growth and development rely on the absorption of water and nutrients in the root. In vascular plants, the absorbed solute in roots is transported through the xylem to reach arial organs. The transverse path of the mineral nutrients from the soil to xylem is across three different type of tissues: dermal, ground and vascular tissue. Endodermis, the innermost layer of the ground tissue, forms a hydrophobic cell wall modification, called the Casparian strip (CS) (1, 2). This ring-like hydrophobic structure not only provides the resistance to negative water pressure, but allows selective transport across the endodermal cell layer, and thus is often regarded as plant inner skin (3–5). The apoplastic continuity of the root is disrupted by the Casparian strip of endodermal cells (6).

It has been long thought that CS is a suberin-based structure (7–10). But a recent study using genetic and pharmacological interferences of lignin and suberin production indicates that the functional CS is made of a lignin polymer (10). The formation of CS starts in around 12^th^ endodermal cells after the onset of cell longitudinal expansion in Arabidopsis roots (11). Recently, a series of breakthrough was made which greatly enhanced our understanding of the molecular components behind CS formation in plants (12–23). CASPs (Casparian strip membrane domain proteins), a family of transmembrane proteins, were discovered to be indispensable for the localized deposition of CS (12). CASP1 localizes specifically to the central zone of endodermal cells, preceding the formation of functional CS (12). Therefore, CASPs were proposed to form a transmembrane scaffold that can assembly lignin biosynthetic machinery, consisting of Respiratory Burst Oxidase Homolog F (RBOHF), Peroxidase 64, Enhanced Suberin 1 (ESB1), and possibly a few other uncharacterized factors (13). The restricted action of these factors in CS membrane domain relies on the CASPs localization, which requires the combinatorial function of the receptor-like cytoplasmic kinase SGN1 and SGN3 (18, 19).

The initial specification of endodermal cells in meristem is under the control of the SHR and the SCARECROW (SCR) transcription factors (24, 25). Lack of SHR leads to the entire loss of endodermal layer and results in a single ground tissue layer with cortex cell identity (24). SCR, a SHR direct target, was proposed to regulate the transition from proliferation to differentiation in the endodermis by activating MYB36 expression (17). Interestingly, *MYB36* loss-of-function mutant exhibits delayed formation of functional CS in the root (16). Genome-wide transcriptome analysis showed that a set of genes regulating CS formation, including CASP genes, ESB-like genes, and *PER64* were all downregulated in *myb36-1* (16).

However, our understanding of SHR pathway and its roles have been mostly restricted to the CEID and its derivative endodermis in the meristem (26, 27). The activation of MYB36 by SHR/SCR was mostly based on the expression analysis in the mutant backgrounds (17). Thus it is possible that SHR solely functions to specify the endodermis in the meristem. Recent studies suggested there might exist spatial restriction of SHR ability to induce endodermal differentiation (28–31). Expressed under the AtSHR promoter in the stele, SHR rice homologues moved beyond endodermal cell layer. Although extra ground tissue layers were observed, CS-containing endodermis was only detected in the cell layer directly contacting the stele (31, 32). Thus, the lack of spatiotemporal resolution of SHR function prevents us from understanding what exact roles SHR plays in the endodermal maturation.

Here, we systemically examined the SHR functions in endodermal differentiation and tried to utilize the synthetic approach to reconstruct fully functional Casparian strips in plant roots. Our results revealed that SHR is essential for defining endodermis at all developmental stages. Transported symplastically from the stele, SHR activates a hierarchical cascade for CS formation, in which MYB36 presumably serves as a relay hub to promote the expression of CASPs, PER64 and ESBs. Beyond that, SHR turns on the specific expression of SGN3, a kinase regulating the CASPs and CS positioning, and this is independent of MYB36. SHR direct target, SCR is dispensable for the activation of these components. However, SCR appears to participate in positioning CASPs to the CS zone of the endodermal cell membrane. Despite of this hierarchical network, SHR mediated pathway needs stele-derived peptides to sufficiently establish an intact CASPs band and CS. Thus, our results provide a genetic framework for endodermal differentiation and reveal a unique non-cell-autonomous mechanism by which the combination of two parallel signals (in both symplastic and apoplastic manners) derived from the stele promote the establishment of functional CS in the neighborly contacting tissue.

## Results

### 1. SHR and SCR distinctly regulate the casparian strip formation

SHR/SCR pathway specifies endodermal cell fate in root apical meristem. CS is a specialized cell wall modification in differentiated endodermis. It is possible that impaired endodermal specification in the meristem can block CS formation in the endodermis. We tested this by examining the presence of the functional CS using Propidium iodide (PI) exclusion assay (10). As expected, *shr-2* exhibited the penetration of PI into the stele all along the root, indicating the loss of the entire CS. However, *scr-4* only had a delayed PI penetration but still blocked the PI in upper region of the root. Compared to wild-type (WT), in which PI exclusion occurred around 4-5 cells after the initial rapid cell elongation, *scr-4* blocked the PI much later (in round 24–25th cells) (supple Fig. 1). This indicated that SHR is indispensable for CS formation, but SCR possibly regulates the temporal formation of the CS.

Prior to the CS formation, CASPs localize to the central zone of endodermal cell membrane and this was proposed to serve as a scaffold to assemble CS catalyzing enzymes. We therefore visualized the pCASP1:CASP1-GFP in both *shr-2* and *scr-4*. No CASP1-GFP was seen in *shr-2* mutant, while CASP1-GFP can be clearly observed in the mutant cell layer of *scr-4* root (Figure 1a). But surprisingly, closer observation showed that unlike the normal central zone localization, CASP1-GFP moved to the stele-facing corner of the mutant cell layer in *scr-4* (Figure 1a-e).

**Figure 1.**
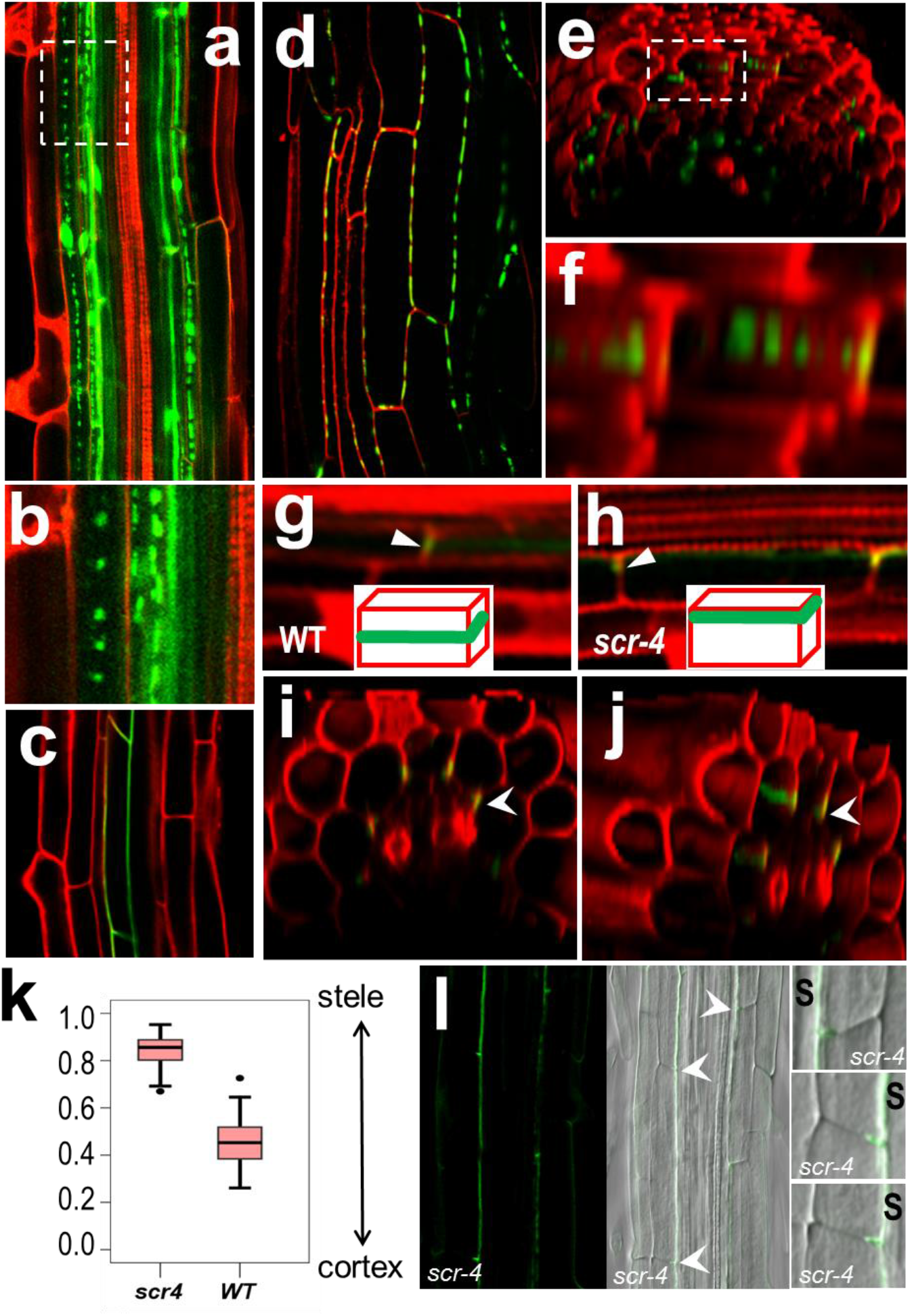
SHR and SCR have distinct roles in promoting CS. a-f) The expression of pCASP1:CASP1-GFP in *35S:SHR-GFP* (a, b), wild type (c) and *pG1090-XVE::SHR* (d-f) roots. *pG1090-XVE::SHR* roots were treated by estradiol for 2 days. (b) is zoomed view of the boxed region of (a) in which the beadstring-like structures of CASP1-GFP are clearly visible. (e & f) 3-D reconstruction of *pCASP1:CASP1-GFP* in *pG1090-XVE::SHR* roots. (f) is the zoomed view of the boxed region in (e). g-j) The expression of *pCASP1:CASP1-GFP* in wild type (g) and scr-4 (h-j) roots. White arrow heads and cartoons in (g & h) show the relative position of CASP1-GFP. Cross-section (i) and 3-D view (j) of *pCASP1:CASP1-GFP* clearly show the stele-facing localization of CASP1-GFP (pointed by white arrow heads) in the ground tissue layer of *scr-4*. k) Quantification of CASP1-GFP positions shown in (g-j). Y axis represents the relative position of CASP1-GFP (%) with stele at 1.0 and cortex at 0.0 (n ≥ 24 cells). i) Lignin (green) deposition in longitudinal section of *scr-4*. Arrowheads point to the lignified position of mutant cell layer that is enlarged in right.

It has been demonstrated that CS is mainly made of lignin (10). Although PI exclusion and CASP localization can reflect the presence of functional CS, a more direct way is to detect the lignified band on the cell wall (Alassimone et al., 2010). In WT, two autofluorescence dots of lignified CS band can be readily seen in the apical/basal cell wall of endodermis. This became invisible in *shr-2*, indicating the loss of CS (supple Fig. 2). Consistent with mis-localization of CASP1, the lignified CS band also shifted to stele-facing corner of the mutant cell layer in *scr-4* root (Figure 2F).

**Figure 2.**
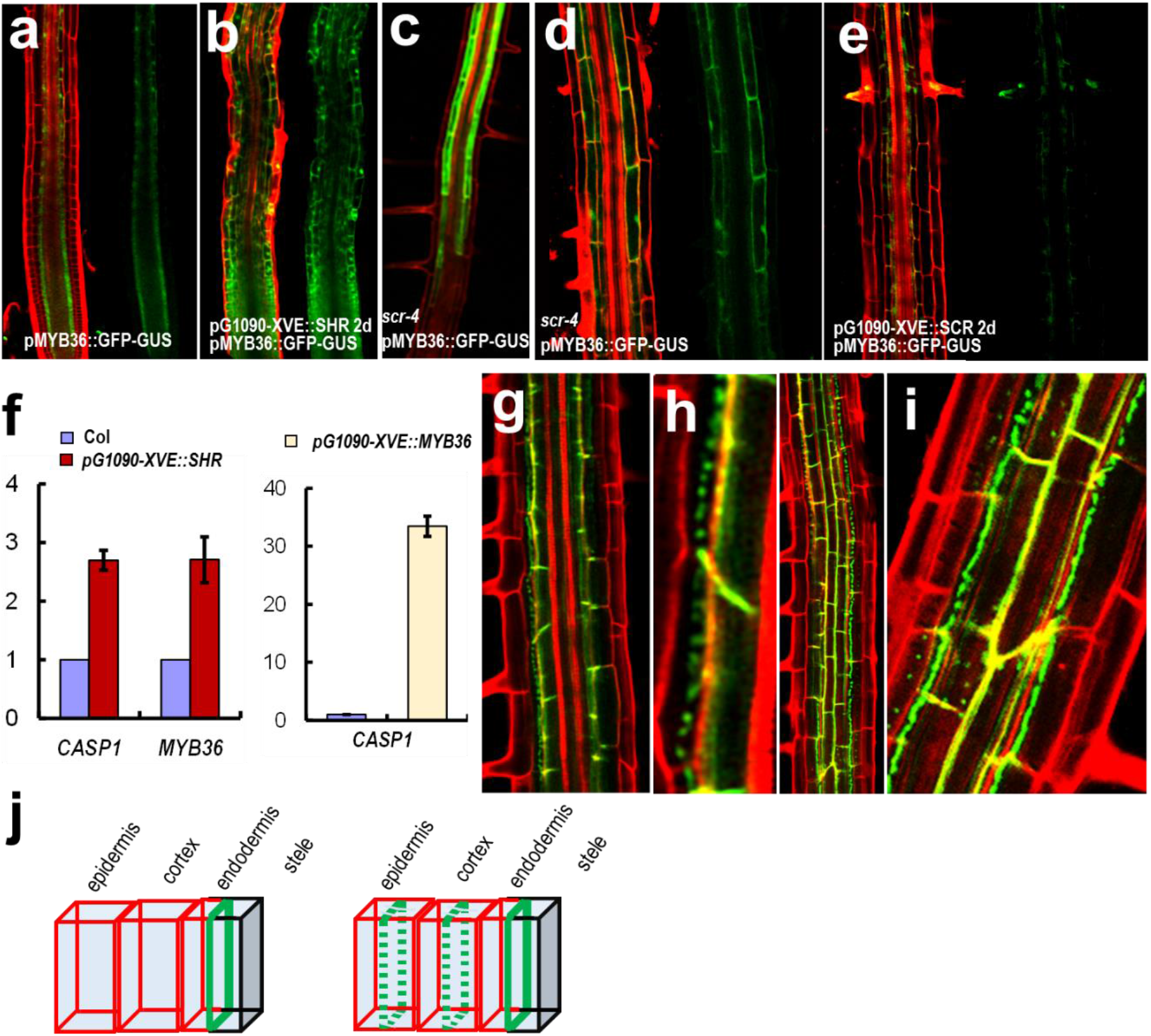
MYB36 activates *CASP1* expression but is not sufficient for forming intact CASP1-GFP bands. a-e) The expression of *pMYB36:GFP-GUS* in wild type (a), *pG1090-XVE::SHR* (b), *scr-4* (c, d) and *pG1090-XVE::SCR* (e) roots. Both *pG1090-XVE::SHR* and *pG1090-XVE::SCR* roots were treated by estradiol for 2 days. f) Activation of *CASP1* and *MYB36* by SHR, and *CASP1* by MYB36 using qRT-PCR. Y axes represent the relative expression of the transcripts. g-i)) The expression of *pCASP1:CASP1-GFP* in *pG1090-XVE::MYB36* roots that were treated by estradiol for 2 days. Note the discontinuous beadstring-like structures that are similar to CASP1-GFP in SHR-overexpressing roots. j) Cartoon depicting the PI staining (in red), non-PI staining (in black) and CASP1-GFP (in green) in wild type roots (left) and *pG1090-XVE::MYB36* roots that were treated by estradiol for 2 days (right). Different cell layers are marked.

### 2. SHR promotes patchy CASP1 distribution via MYB36

Although SHR is indispensable, it is not clear whether SHR mediated pathway is sufficient to promote CS. To address this, we ectopically expressed SHR in Arabidopsis roots. SHR activated CASP1 expression in almost all cell types except the epidermis. Despite the expression, CASP1-GFP appeared to form discrete punctae and aggregation in extra ground tissues (Figure 3b-d). In many cells, the patchy CASP1 distribution appeared to align correctly in the CS zone of the cell wall (Figure 3b&d).

**Figure 3.**
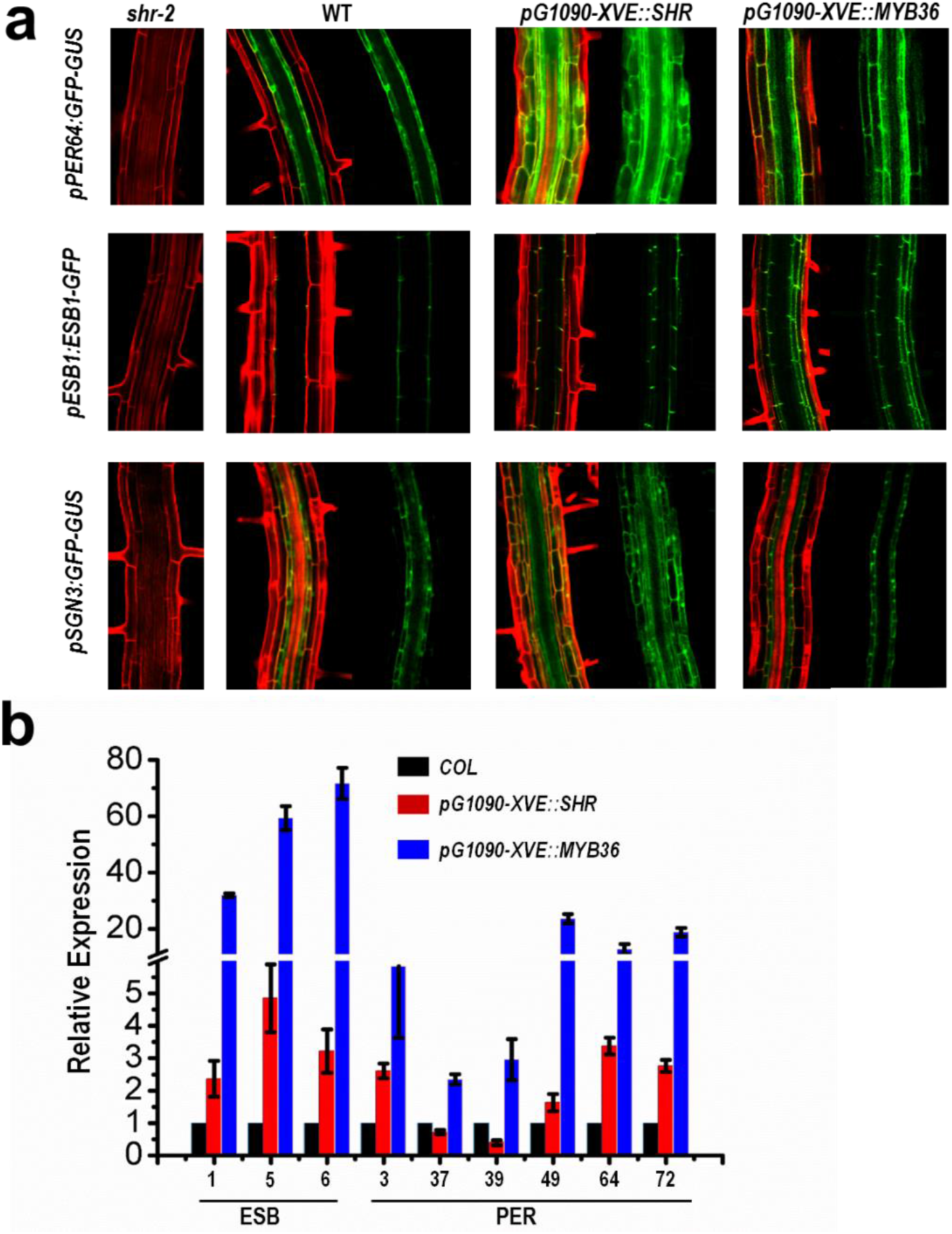
SHR and MYB36 indirectly activate essential genes for CS. a) The expression of *pPER64:GFP-GUS*, *pESB1:ESB1-GFP* and *pSGN3:GFP-GUS* in shr-2, wild type, *pG1090-XVE::SHR*, and *pG1090-XVE::MYB36* roots. Both *pG1090-XVE::SHR* and *pG1090-XVE::MYB36* roots were treated by estradiol for 2 days. Note the SGN3 can be activated by SHR but not by MYB36. b) Activation of *ESB* and *PER* genes (X axis represents the gene number of each homologous member) by both SHR and MYB36 using qRT-PCR. Note the stronger activation by MYB36 than by SHR.

**Figure 4.**
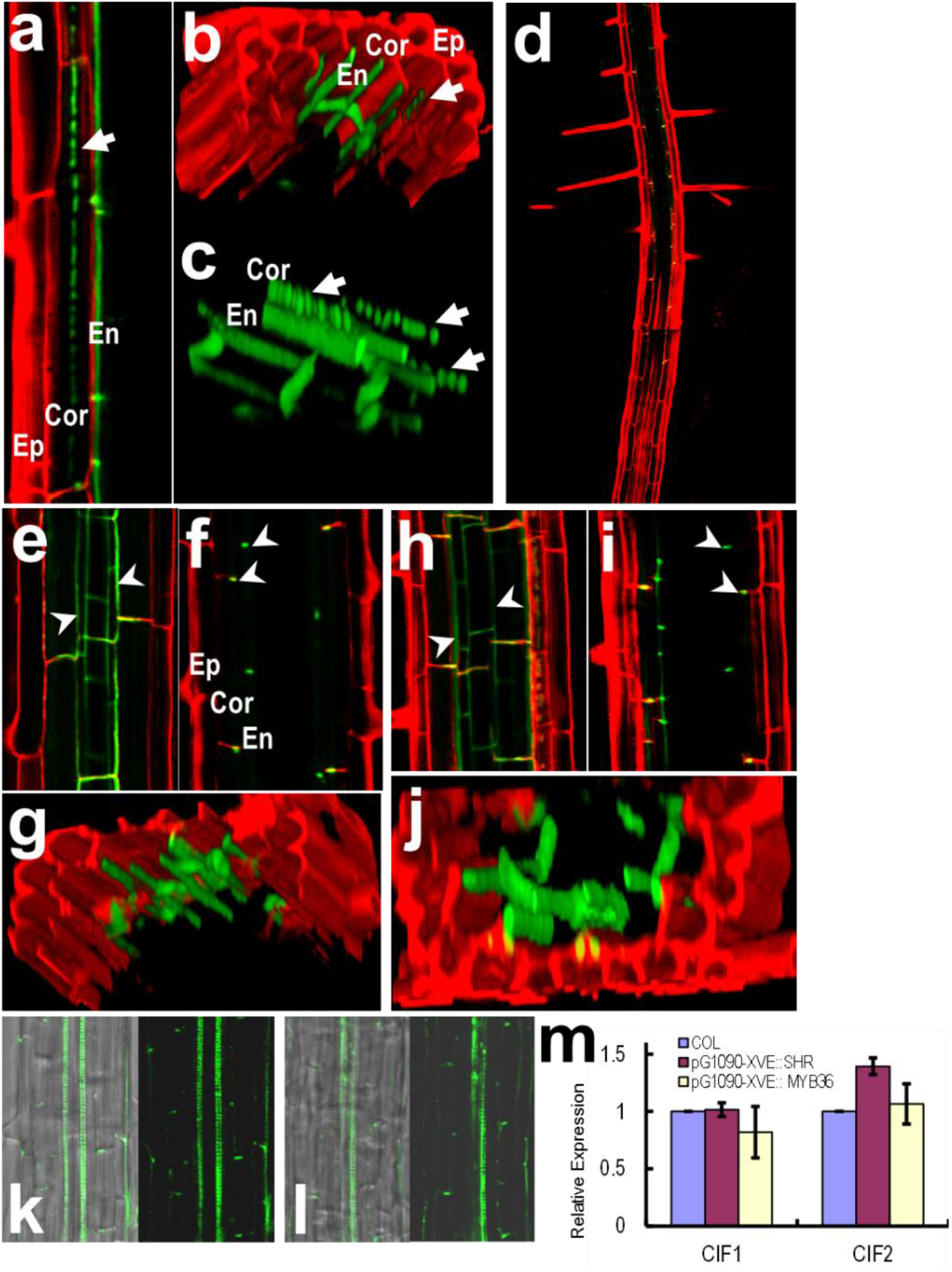
SHR and small peptides CIF1/2 operate in conjunction to promote intact CASP1 and CS in cells outside of endodermis. a-c) The expression of *pCASP1:CASP1-GFP* in *pG1090-XVE::SHR* (treated by estradiol for 2 days) roots. Ep, epidermis; Cor, cortex; En, endodermis. d-j) The expression of *pCASP1:CASP1-GFP* in *pG1090-XVE::SHR* roots treated by estradiol together with CIF1 (d-g) or CIF2 (h-j) for 2 days. (d) shows the penetration of PI only in epidermis. (e & h) show the intact CASP1-GFP in multiple cell layers (pointed by arrow heads). (f & i) show the dots of CASP1-GFP bands on apical and basal sides of the cells (pointed by arrow heads). (g & j) show the continuous CASP1-GFP localization in cells outside of endodermis which is different from the discontinuous beadstring-like structures in (b & c). (k & l) Lignin (green) deposition in *pG1090-XVE::SHR* roots treated by estradiol together with CIF1 (k) or CIF2 (l) for 2 days. m) Transcriptional level of CIF1 and CIF2 was not activated by SHR and MYB36, shown by qRT-PCR.

**Figure 5.**
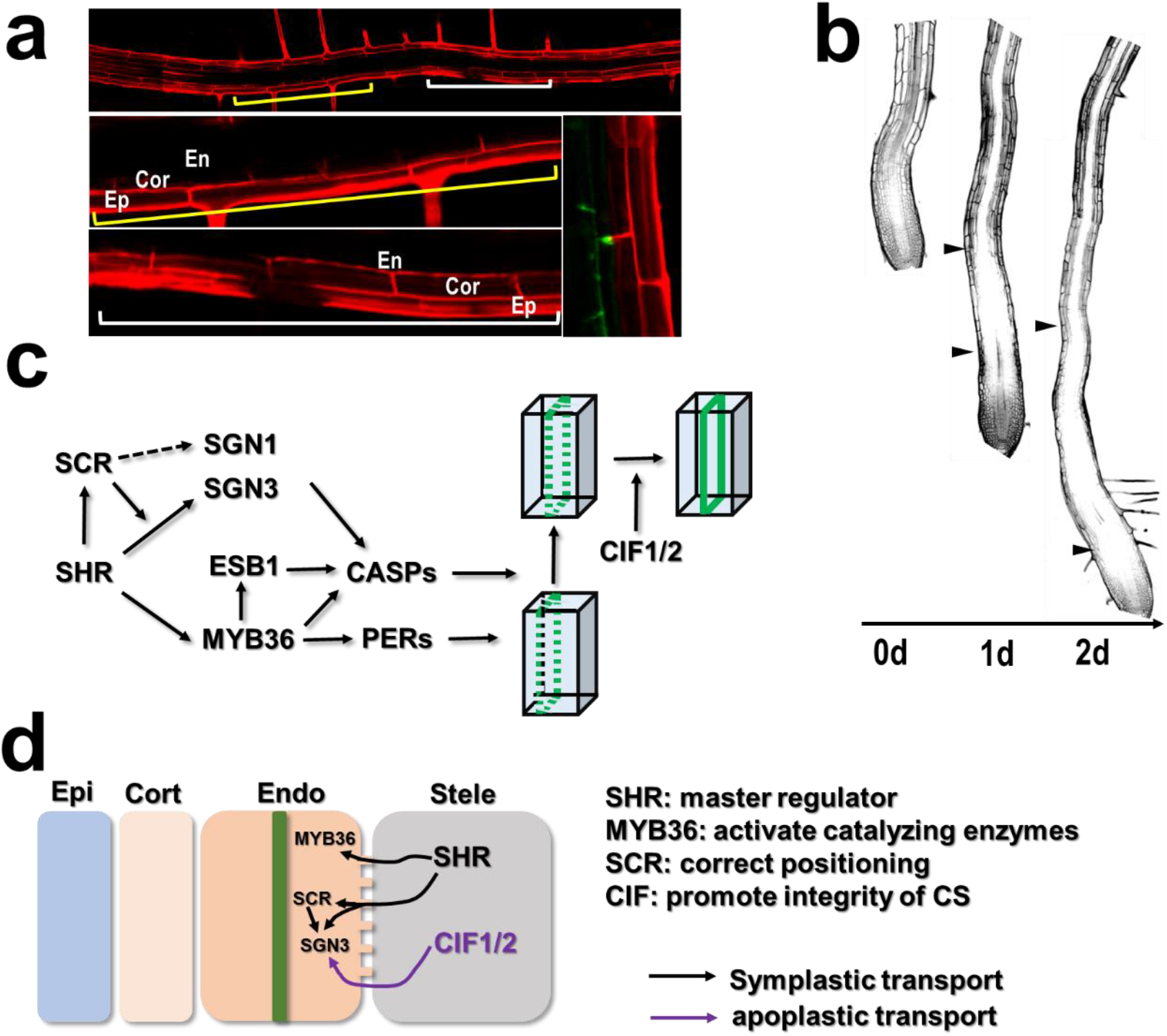
Stele derived signals promote CS formation in a restricted zone. a, b) PI exclusion assay in *pG1090-XVE::SHR* roots treated by estradiol together with CIF1 for 2 days (a) and *35S:SHR-GFP* roots treated by CIF1 for 1-2 days (as labeled) (b). Arrowheads in (b) point to the start and end of the PI exclusion zone. Note the expansion of PI exclusion zone in the 2d treatment. c) Schematic of possible SHR cascades during CS formation. Green line and dotted green lines represent intact CASP1 and discontinuous CASP1 respectively. d) The putative working model for the formation of functional CS. Two independent stele-derived signals promote CS formation via symplastic (in black) and apoplastic (in purple) transport. Green band represents intact CS formed in endodermis.

To rule out the artifacts possibly caused by SHR overexpression, we examined CASP1-GFP in estradiol inducible *pG1090-XVE::SHR* roots. Instead of forming the ring-like structure, CASP1-GFP in *G1090:XVE-SHR* roots also exhibited discrete punctae along the cell membrane in extra ground tissues (Figure 3f). The 3-D reconstruction showed that CASP1-GFP punctae aligned beneath the cell wall in *pG1090-XVE::SHR* expressing roots (Figure 3h&i).

It was recently shown that the formation of functional CS was delayed in MYB36 loss-of-function mutant (16, 17). Consistently with this, a set of CS genes including CASPs, ESBs and PER64 were all downregulated in *myb36-1* (16). Interestingly, expressing SHR and SCR in respective mutant backgrounds enhanced MYB36 expression (17). Here we tested whether SHR is able to trigger MYB36 expression in cells outside of endodermis. In *shr-2* roots, no *pMYB36:GFP-GUS* expression was seen. But in the *pG1090-XVE::SHR* expressing roots, *pMYB36:GFP-GUS* was activated in all cell types, even including epidermis. Interestingly, in *scr-4*, *MYB36* expression was not abolished, indicating SHR rather than SCR is sufficient for the activation of *MYB36*.

We further examined the CASP1-GFP with *MYB36* expression in different cell types. With estradiol treatment, *pG1090-XVE::MYB36* expression triggered ectopic activation of CASP1-GFP in ground tissues. Similar to *SHR*, *MYB36* expression in non-endodermal cells resulted in discontinued punctae of CASP1-GFP, suggesting *SHR* and *MYB36* presumably function in the same pathway in controlling CASP1 (supple Fig. 3). To verify the possibility that SHR functions via MYB36, we examined CASP1-GFP in *shr-2* roots with the expression of *pG1090-XVE::MYB36*. After the estradiol induction, we clearly saw that *MYB36* recued the expression of CASP1 in the mutant cell layer of *shr-2* (Figure 3j). However, CASP1-GFP accumulated at the stele-facing corner of the mutant cell layer, indicating the CASP1 positioning mechanism is independent of *MYB36* (Figure 3k&l).

### 3. SHR is the master regulator of CS lignification and positioning

In the current model, locally positioned CASPs provide the scaffold for the concerted action of NADPH oxidase and peroxidases. We thus examined whether SHR-SCR-MYB36 circuit regulates those catalyzing enzymes. PER64, the critical enzyme catalyzing the last step from monolignol to lignin in the CS zone (13, 33). We visualized *pPER64:GFP-GUS* reporter in different backgrounds. When SHR was ectopically expressed, *pPER64:GFP-GUS* activation was observed in almost all cell types. In *scr-4*, however, *pPER64:GFP-GUS* had the similar expression pattern as in WT, suggesting SHR, rather than SCR controls *PER64* expression. Similar to *CASP1*, *PER64* can also be activated by *MYB36* in most cell types. Recently, peroxidase-mediated polymerization of lignin has been shown to rely on localized H_2_O_2_ production catalyzed by NADPH Oxidase. A NADPH oxidase, Respiratory burst oxidase homolog F (RBOHF)/SGN4 controls this superoxide local production in CS zone (13). qRT-PCR, however, showed that SGN4 was not upregulated by either SHR or MYB36. This result is consistent with the constitutive expression pattern of SGN4 in roots, and suggests that SHR/MYB36 presumably only activate those genes that are specifically expressed in endodermis.

In addition to locally catalyzed lignification, CS formation also requires correct positioning. This is under the control of the combination of CASPs, ESBs and SGN1/3 (12–14, 18, 19). In the WT, those genes are only active in the endodermis. When *SHR* or *MYB36* was ectopically expressed, *pESB1:ESB1-GFP* expression can be observed in all ground tissues. Consistently, qRT-PCR indicated that other *ESB* genes including *ESB5* and *ESB6* were also upregulated by both SHR and MYB36. However, SGN3, the receptor-like kinase that is essential for localizing CASPs, appeared to be controlled only by SHR. The expression of *pSGN3:GFP-GUS* in *pG1090-XVE::SHR* expanded to all ground tissues as well as epidermis, while *SGN3* expression in *pG1090-XVE::MYB36* roots resembled that in the WT. In *scr-4*, the expression of *PER64, ESB1* and *SGN3* was detected in the mutant cell layer, indicating the SCR is not essential for activation of those genes.

In order to understand the CS regulatory cascade mediated by SHR and MYB36, we examined the transcription level of previously described genes that are involved in lignin biosynthesis and deposition (33, 34). ESB5 and ESB6, previously described as down-regulated genes in *myb36*, are activated by both SHR and MYB36 (16). An ATP-binding cassette transporter, AtABCG29, was reported to transport monolignols from the cytosol to the cell wall (34). This gene exhibited upregulation when SHR or MYB36 were inducibly expressed. The catalysis of the last step from monolignol to lignification is mediated by peroxidase (13, 33). In addition to PER64 we examined, there are eight other peroxidases that are specifically expressed in endodermis (13). We analyzed their transcription in *pG1090-XVE::SHR* and *pG1090-XVE::MYB36.* MYB36 activated all peroxidase examined but SHR only promoted the expression of *PER3, PER49, PER64* and *PER72*.

Putting together the fact that MYB36 failed to activate *SGN3*, we proposed that SHR is the master regulator controlling both catalyzing, transporting and positioning steps during CS formation. MYB36 is an intermediate hub in the whole regulatory hierarchy, which presumably functions as a more direct regulator of many downstream components.

### 4. SHR-SCR-MYB36 circuit is not sufficient to establish a functional CS

Since many components regulating CS formation were activated by SHR, we next asked whether SHR mediated pathway is sufficient to induce a functional CS in cells other than endodermis. Expression of *pG1090-XVE::SHR* produced additional cell layers out of the endodermal layer in the Arabidopsis root. However, we only detected a single lignified cell layer surrounding the stele in roots expressing *pG1090-XVE::SHR* even after 2-day estradiol treatment. Consistent with this, we saw the exclusion of PI only from the ground tissue layer immediately surrounding the stele, indicating SHR is not sufficient to produce functional CS in extra cell layers. Similarly, ectopic expression of MYB36 did not create PI proof layer in cells beyond endodermis. The lignin autofluorescence in the root expressing *pG1090-XVE::MYB36* showed the similar pattern to the one expressing SHR. These results indicate that despite of activation of a group of critical components of CS forming machinery, neither SHR nor MYB36 was sufficient to establish the functional CS.

### 5. CIF1/2 operate parallel with SHR to promote fully functional CS

In all cases above we observed, only the cell layer that is directly contacting stele acquire the ability to form function CS, implying that additional stele derived signals are indispensable. Recently, two stele expressed small peptides, CIF1 and CIF2, were identified that bind to the SGN3 receptor to form intact CS (20, 21). Compared to the discontinuous punctae of CASP1-GFP in the control of mock treatment, addition of CIF1 or CIF2 induced continuous CASP1-GFP ring in cells outside of endodermis. Except for the epidermis, all ground tissues showed two CASP1-GFP dots at the correct CS zone. In line with this, lignin autofluorescence staining exhibited additional lignin deposition in non-endodermal cells. Using PI penetration assay, we found that PI was blocked immediately after epidermis, indicating the expansion of functional CS to the outmost ground tissue layer. The expression of *CIF1* and *CIF2* seemed to be independent of SHR/MYB36 as the transcriptional level of CIFs remained unchanged when *SHR* or *MYB36* was ectopically expressed.

Interestingly, the cooperative operation of SHR and CIFs seemed to have zone restriction. The functionality of the combination of SHR and CIFs only occurred in the region that is immediately above the expansion zone. Compared to 1d treatment of CIFs, *35S:SHR-GFP* roots showed more expanded PI exclusion zone (excluded from all ground tissues) if treated with CIFs for 2d.

In summary, we propose the model in which two independent stele-derived signals promote CS formation via both symplastic and apoplastic transport. Through activating *MYB36*, SHR promotes the expression of a number of critical enzymes and genes including CASPs, ESBs and PERs. On top of this, SHR also activates kinases that help correctly position the CS, including SGN1 and SGN3. This cascade work together with a parallel apoplastic signal mediated by stele-derived peptides, CIF1 and CIF2, to finalize the CS establishment in roots.

## DISCUSSION

Plant cells are immobile within a tissue and the cell fate determination thus mainly relies on the position of each individual cell. To provide position-dependent cell fate regulation, plant cells need to communicate via the signaling over tissue boundaries. Nearly all plant cells are connected by plasmodesmata, which underscores the importance of symplastic communication between plant cells (35). The initial endodermal specification has been a typical example of coordinated asymmetric cell division and cell fate determination regulated by the mobile regulator SHR (36, 37). Our understanding of SHR functions in endodermal specification mostly derived from the studies in CEID and its derivative immature endodermis (26, 27). In this study, we presented the evidence indicating that SHR moves symplastically across the tissue to mediate a hierarchy of signaling cascade to promote the CS formation in the mature zone. This signaling cascade appeared to function also in cells beyond endodermis. However, the cells outside of endodermis only formed discontinued CASP1 punctate on the cell wall despite of the expression of a set of CS regulators induced by SHR. This is consistent with the previous report that SHR is necessary but not sufficient for specification of a functional endodermis (31, 32). Interestingly, when we added the stele-derived small peptides, CIF1 and CIF2, into this inductive system mediated by SHR, we were able to create intact CASP1 structures and the functional CS in cells beyond the endodermis. CIFs, although expressed in stele, are not under control of SHR. Thus, this is a demonstration of the deployment of two parallel signaling systems by plants for coordinately specifying the cell fate.

In animals, mobile signals such as Hedgehog and wingless can travel between tissues to direct their boundaries (38). Those regulators in animals mostly are secreted outside of the cell to transduce the short-range signaling (39). In plants, however, signaling molecules transport from cell to cell not only through apoplastic secretion, but via symplastic communication mediated by plasmodesmata (37). CS formation represents the elaborate control of plant development by joined force of two independent signaling systems, in which both apoplastic and symplastic communication were employed. Our results also explain why although essential, SHR alone is not sufficient to confer endodermal cell fate in ground tissues. In this regulatory system, SHR is restricted to the endodermis by its downstream target SCR and CIF1/2 are blocked by the intact CS, indicating that plants exploit these two locking mechanisms to ensure only single layer of ground tissue directly contacting stele acquire all necessary components for CS formation. One possible advantage of this two-way locking strategy is that either further movement of SHR or CIFs is not able to induce functional CS. The response to the SHR induction is thus confined to the endodermis and this also creates a clear-cut boundary, rather than a transition zone caused by the effect of morphogen gradients of CIF1/2. Through the coordinative action of a mobile transcription factor and mobile small peptides, plant roots thus can precisely create a boundary between the vascular tissue and the ground tissue.

The regulation mediated by SHR and CIF1/2 seemed to be sufficient to rebuild CS in cells outside of endodermis. However, this regulation longitudinally functioned in a restricted area, which locates above the elongation zone. Thus the endodermal maturation appeared to be an assembly line with CS formed only in one of processes. In addition, we rarely observed CASP1-GFP induced by SHR in epidermis, suggesting a possible inhibitory mechanism specific to epidermis blocks SHR mediated signaling cascade. Therefore, CS forming machinery can also be fine-tuned by other developmental programs regardless of its sufficiency in establishing CS.

In *shr-2*, CS was entirely abolished while CS formation either delayed or lost integrity in the mutants of other CS forming regulators. In the regulatory cascade of CS formation, SHR likely serves as a master regulator and an upstream hub of the whole hierarchy. In our study, SHR not only activated *MYB36*, the sub-hub regulator promoting the expression of a number of critical enzymes, but also activated the components controlling the correct localization of the CS. MYB36 acts as a core part of the cascade downstream of SHR, but SGN3 is under the control of SHR rather than MYB36. Compared to SHR, MYB36 seemed to be a more effective activator of many CS forming components. This is consistent with the fact that the promoter of those components is the target of MYB36 rather than SHR. But interestingly, SHR or SCR did not bind to MYB36 promoter, indicating the complexity of the regulatory hierarchy and the existence of additional layers of regulations. This complex hierarchical cascade could allow higher flexibility of the regulatory system of CS. It would be interesting to know how SHR, as the master regulator of this hierarchy, responds to the environmental and endogenous cues.

### Materials and Methods

#### Plant material and growth conditions

All plants are in the Columbia (Col) accession. The mutants *shr-2* and *scr-4*, and transgenic lines (for details of plant lines and plasmid construction, see Methods) are used. For all experiments, seeds were sterilized, sown on a solid medium containing 0.5×Murashige–Skoog (MS) medium, incubated 2d at 4°C,and grown at 22 °C under a 16-h light/8-h dark photoperiod. For PI blockage, lignin autofluorescence, Fluorol yellow staining, FM4-64 staining, FDA penetration, promoter analysis, and statistical analysis, seedlings were 6 days old at the point of analysis.

#### Molecular cloning and Transformation

For cloning and generation of expression constructs, Gateway Cloning Technology (TRANS) and standard molecular biology techniques were used. The pG109 0-XVE::SHR; pCASP1::CASP1-GFP line was obtained by crossing pCASP1::C ASP1-GFP to pG1090-XVE::SHR. The promoter reporters pPER64 (1478bp bef ore ATG)::GFP-GUS, pMYB36 (2987bp before ATG)::GFP-GUS and pSGN3 (2 522bp before ATG)::GFP-GUS were cloned into the pGWB632 expression vector. Primer sequences are as follows: pPER64-F/R: GGGGTACCTCGTACAATAGCACGGGTCATCACT/CCGCTCGAGCTTTAACAAACTTTTCGAAATTTTAATT; pMYB36-F/R:GGGGTACCCCCACCTCTCAAAACAATAAAT/CCGCTCGAGC TTGTCGTTGTTGTTCTCTTCC; pSGN3-F/R:GGGGTACCCTGAGTGAGATTCATACTTGGTGC/CCGCTCGAGGTTTTCTTCTTCGTCGTCTTATG. The gene cod ing sequences (cDNA) of MYB36(AT5G57620), obtained from The Arabidopsis Information Resource(TAIR), was cloned using Gateway into the PMDC7 expression vector with primers as F/R: AAAAAGCAGGCTTCATGGGAAGAGCTC CATGC/AGAAAGCTGGGTTAACACTGTGGTAGCTCATCTGA.The plasmids w ere transformed in the wide type and *pG1090-XVE::SHR* using an Agrobacterium tumefaciens strain GV3101 by the floral dip method.

#### qRT-PCR

Total RNA was extracted from whole roots using TRIzol reagent (ambion). An equal amount of total RNA (1ug) was used to generate cDNA with TransScript All-in-One First-Strand cDNA Synthesis SuperMix for qPCR (TRANS) and diluted 10-fold for assays. qRT-PCR was performed using TransStart Top Green qPCR SuperMix (TRANS).Three biological replicates performed in three technical replicates were analyzed for each experiment. Primers were designed and tested for amplification efficiency by DNA melting curve analysis and gel electrophoresis of the amplified products. Standard curves were run for each primer pair. Values reported are the means of three biological replicates after efficiency corrected quantification. Relative transcript levels were calculated with eiF4α as the reference gene. The primers were used. (See supplemental table 1)

#### Microscopy and histological analysis

Confocal images were performed on a Zeiss LSM880. Excitation and detection parameters were set as follows: GFP 488nm, 493-548nm; PI 561nm, 566-718nm. For FM4-64 staining, Seedlings were incubated in FM4-64(10uM) for 5min, rinsed and observed. For lignin autofluorescence staining, excitation and emission frequencies for GFP (488nm, 493-548nm) was used. Observation of PI uptake was performed in (11). Lignin autofluorescence was visualized according to (11). Briefly, seedlings were incubated in 0.24 N HCl (in 20% methanol) at 57°C for 15 min, then covered in 7% NaOH (in 60% ethanol) at 25°C for 15 min, in turn, rehydrated in 40%, 20% and 10% ethanol for 5 min, incubated in 5% ethanol and 25% glycerol for 15 minutes and mounted in 50% glycerol for observation. (also see http://wp.unil.ch/geldnerlab/resources-and-protocols/protocols/).

## Acknowledgements

We thank Niko Geldner for materials and comments on the manuscript, Philip Benfey for critically reading of the manuscript, Lei Shi for technical assistance with Zeiss 880 confocal microscope at Cell Biology Core of Fujian Agriculture and Forestry University.

P.L., M.X., Q.Y., B.H. were supported by the 1000-Talents Plan from China for young researchers (Grant KXR15012A) and a start-up fund from Fujian Agriculture and Forestry University awarded to S.W.

## Footnote

The authors declare no competing interests.

---

**Supple figure 1.**
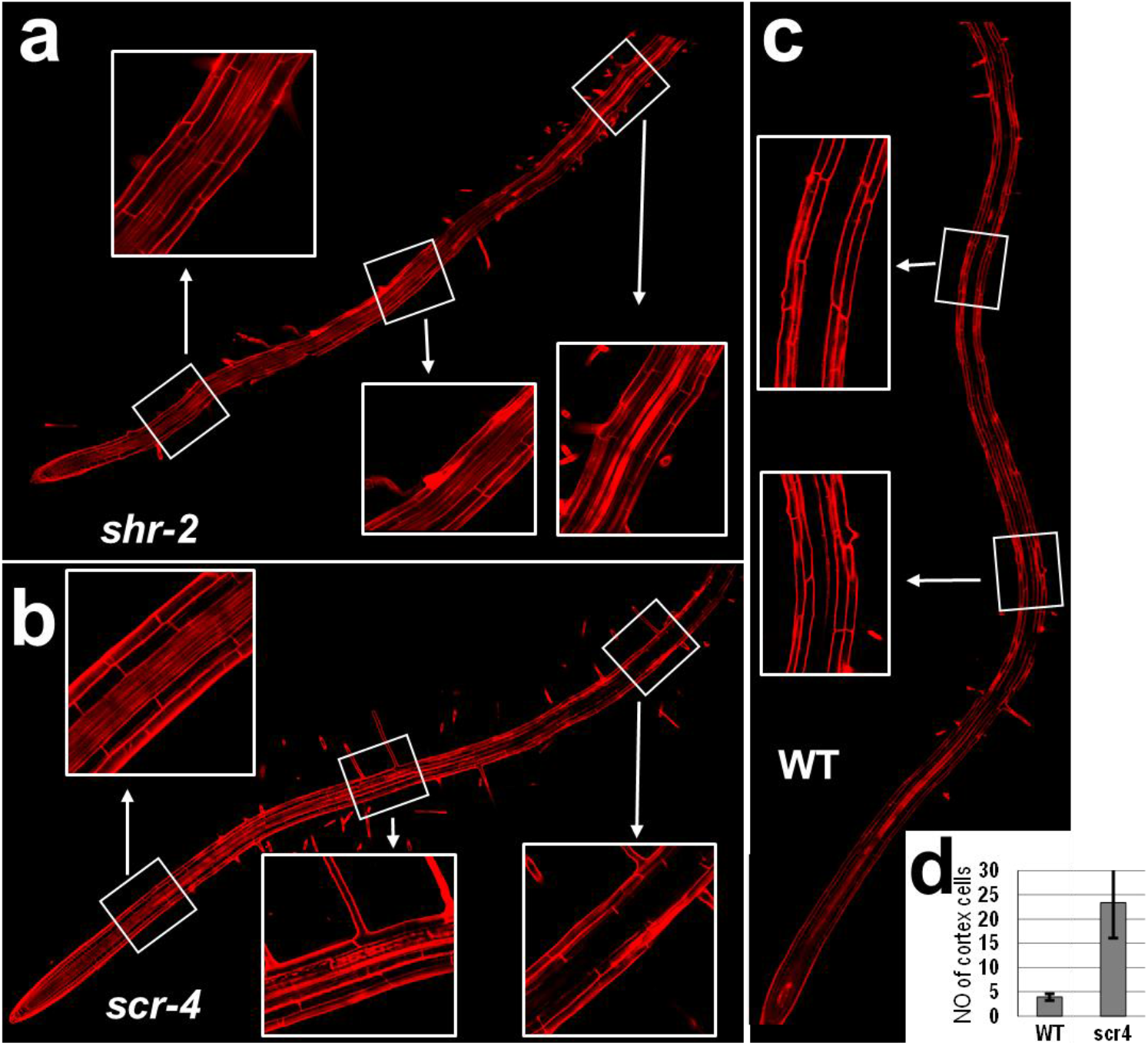
SHR is indispensable for function Casparian strip. PI penetration assay in *shr-2* (a), *scr-4* (b) and wild type (WT) (c). Stitched confocal images (20×) revealing the block of PI along the whole root. The insets show PI penetration across root cell layers in boxed regions. (d) Quantification of PI permeability in WT and *scr-4*. The site of barrier establishment was scored by counting cortex cells after the first root hair. (6-day-old seedlings, mean ± SD,15 < n < 20, Error bars are SD). PI, propidium iodide.

**Supple figure 2.**
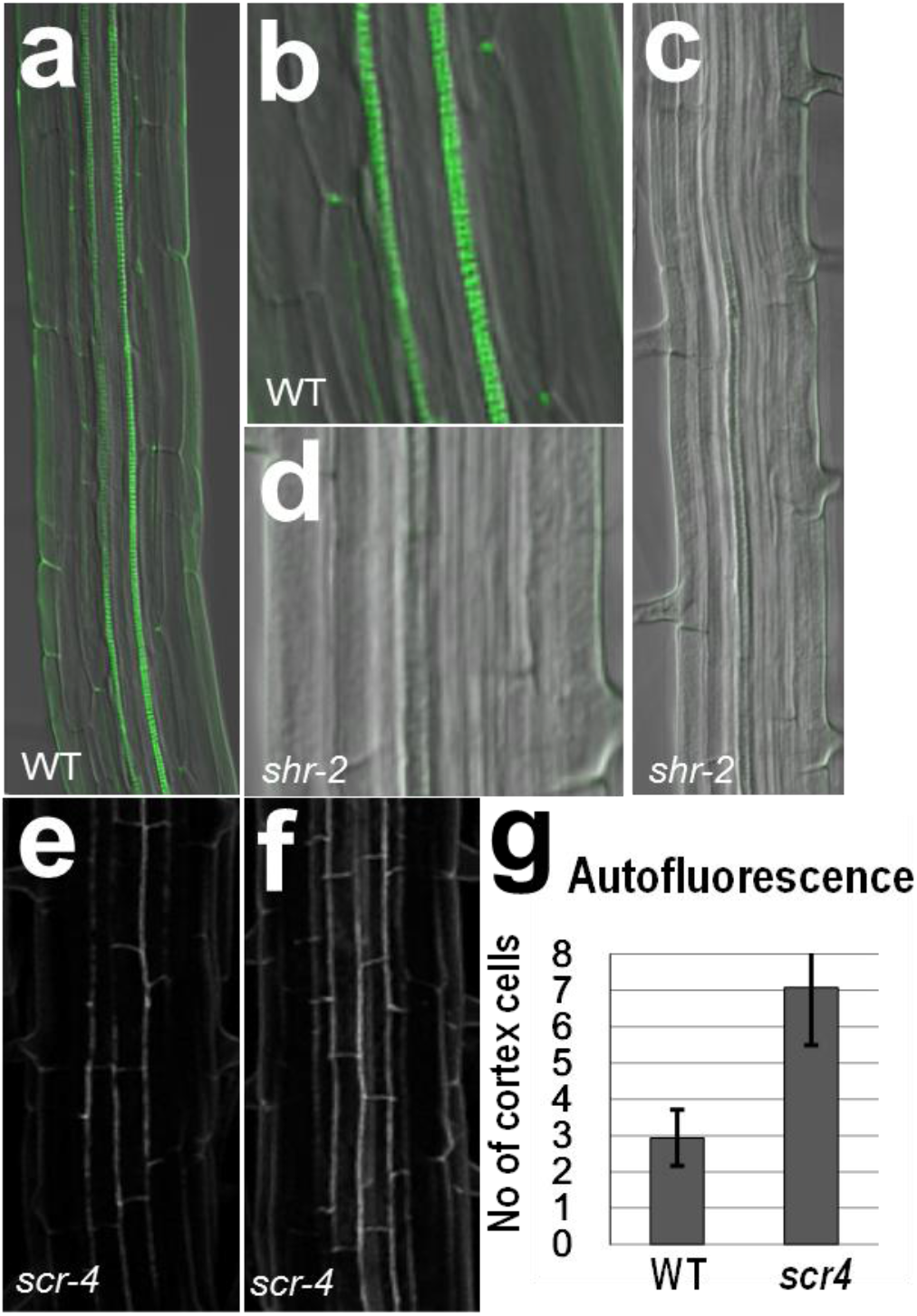
SHR is essential for endodermal lignification. Casparian strip (CS) shown by the lignin autofluorescence staining in WT (a, b) and shr-2 (c, d). (e, f) Z-stack confocal image of the autofluorescence of CS in *scr-4*. Spiral structures in the center of the root are xylems. (e) shows the starting region of CS formation and (f) shows the mature zone. g) Quantification of lignin autofluorescence. Y axis is the initial site of autofluorescence detection scored as the number of cortex cells from the first root hair. (6-day-old seedlings, mean ± SD, n ⩾ 15, Error bars are SD).

**Supple figure 3.**
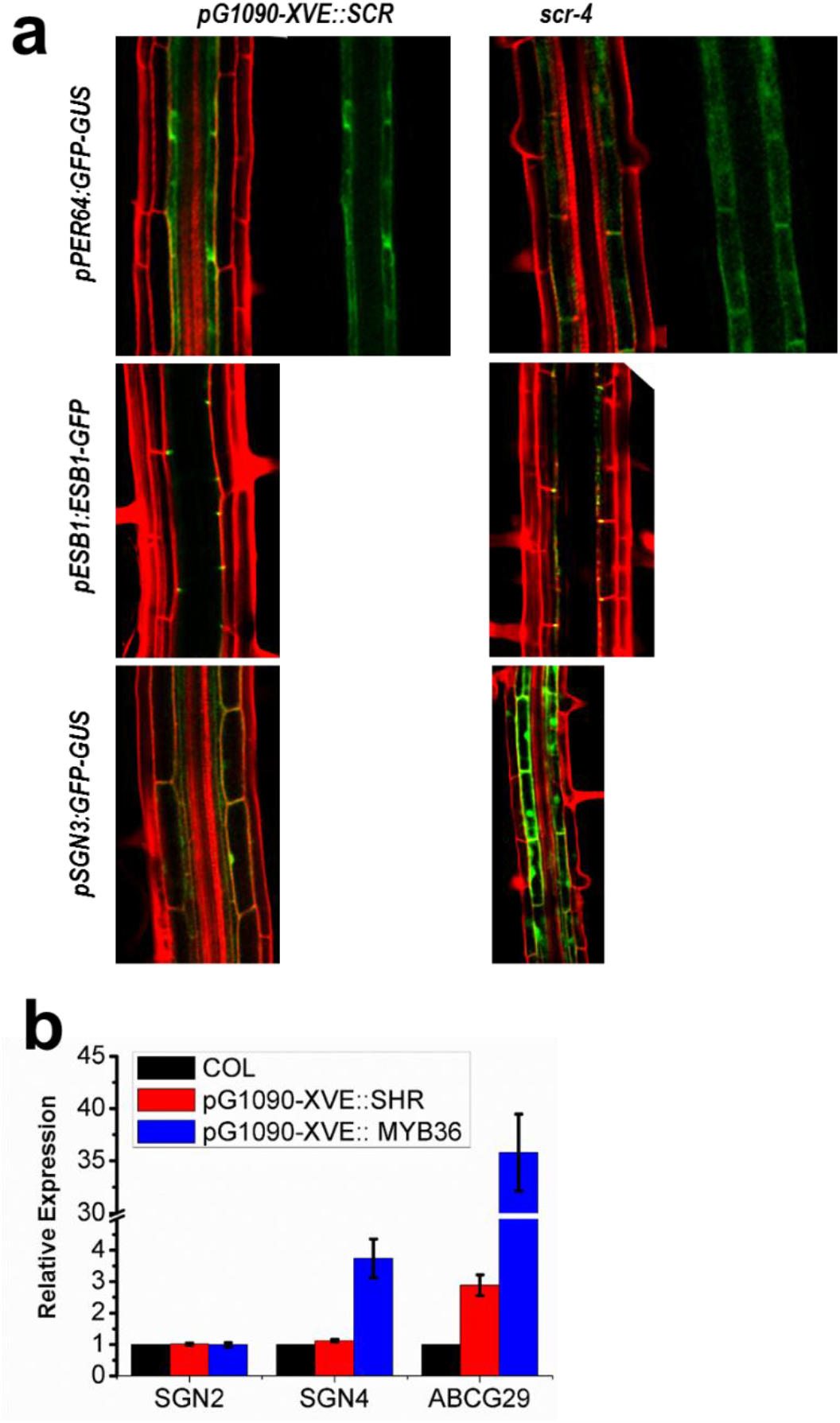
SCR is not essential for the expression of CS genes. a) The expression of *pPER64:GFP-GUS, pESB1:ESB1-GFP* and *pSGN3:GFP-GUS* in *scr-4* and *pG1090-XVE::SCR* (treated by estradiol for 2 days) roots. b) Transcriptional analysis of *SGN2, SGN4* and *ABCG29* with inducibly overexpressed SHR and MYB36.

**Supple figure 4.**
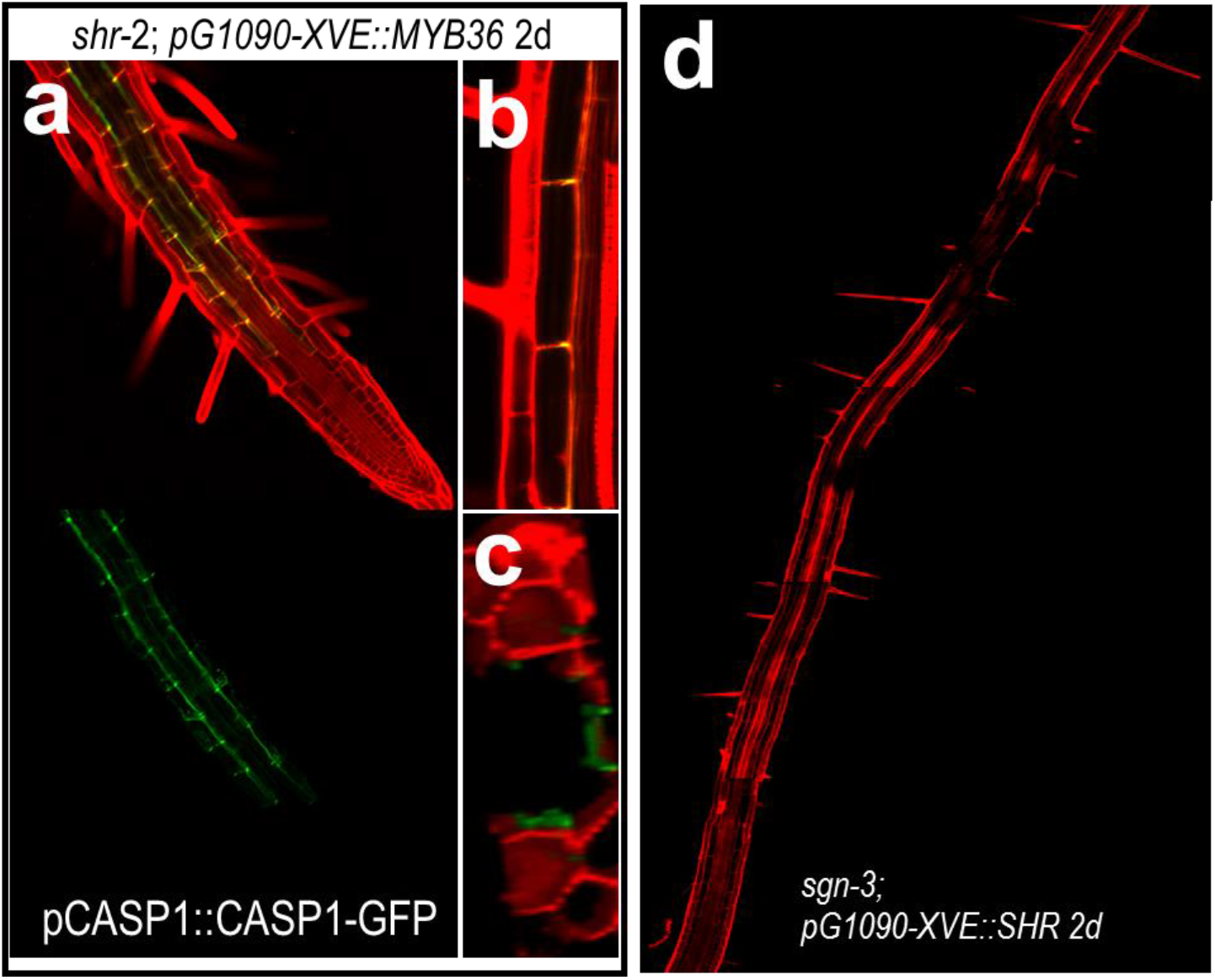
SHR is the upstream regulator of the cascade. a-c) The expression of *pCASP1:CASP1-GFP* in *shr-2* roots with *pG1090-XVE::MYB36* expression (treated by estradiol for 2 days). (b, c) show the stele facing localization of CASP1-GFP in the mutant layer of shr-2. d) PI exclusion assay in the sgn-3 root with *pG1090-XVE::SHR* expression (treated by estradiol for 2 days). Note the expression of SHR did not rescue the delayed PI exclusion in *sgn-3*.

**Supple figure 5.**
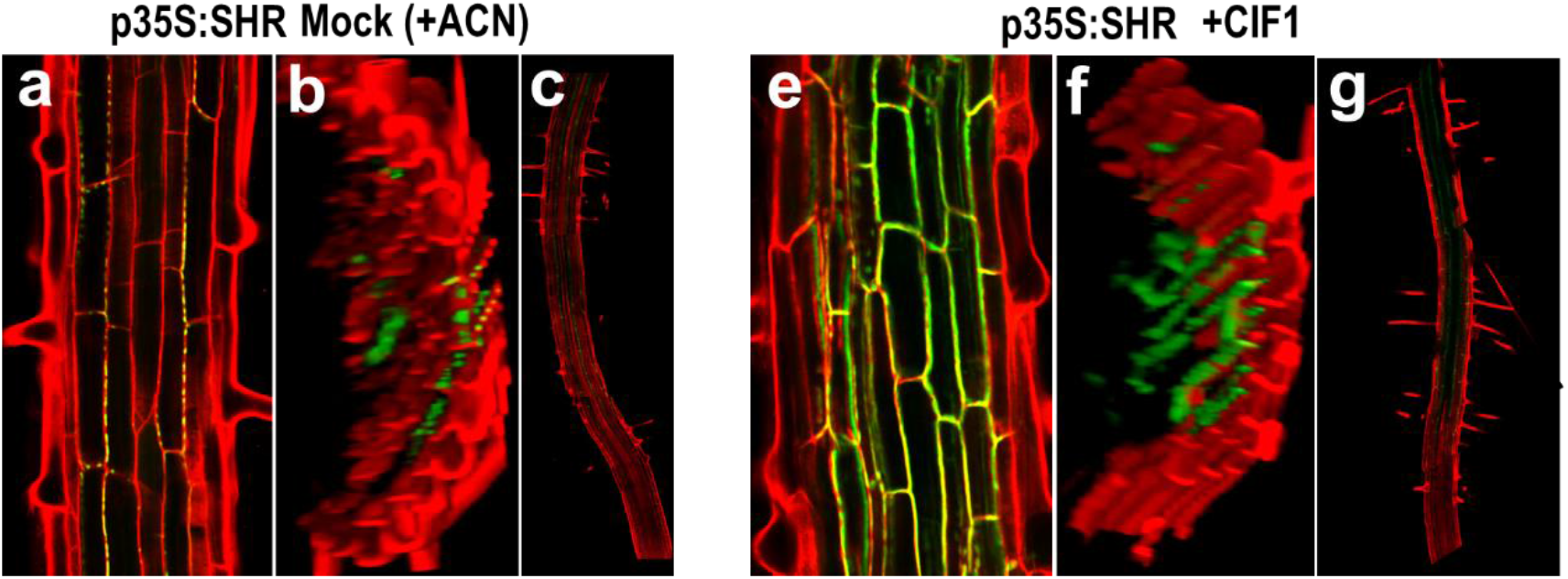
CIF promotes the intact CASP1-GFP bands in non-endodermal cells with ectopic expression of SHR. a-c) The expression of *pCASP1:CASP1-GFP* in *35S:SHR* roots with the treatment of the solvent ACN as the mock. e-g) The expression of *pCASP1:CASP1-GFP* in *35S:SHR* roots with the treatment of the CIF1 for 2 days. (c & g) show the comparison of the PI exclusion.

